# Manipulation of microbiota reveals altered myelination and white matter plasticity in a model of Huntington disease

**DOI:** 10.1101/413112

**Authors:** Carola I. Radulescu, Marta Garcia-Miralles, Harwin Sidik, Costanza Ferrari Bardile, Nur Amirah Binte Mohammad Yusof, Hae Ung Lee, Eliza Xin Pei Ho, Collins Wenhan Chu, Emma Layton, Donovan Low, Paola Florez De Sessions, Sven Pettersson, Florent Ginhoux, Mahmoud A. Pouladi

**Affiliations:** Translational Laboratory in Genetic Medicine (TLGM), Agency for Science, Technology and Research (A*STAR), 138648, Singapore; Department of Psychology, The University of Sheffield, S1 2LT, UK; Lee Kong Chian School of Medicine, Nanyang Technological University, 637551, Singapore; GIS Efficient Rapid Microbial Sequencing, Genome Institute of Singapore, A*STAR, 138672, Singapore; Singapore Immunology Network, A*STAR, 138648, Singapore; Singapore Centre for Environmental Life Sciences Engineering, 60 Nanyang Drive, 637551, Singapore; Department of Medicine, Yong Loo Lin School of Medicine, National University of Singapore, 117597, Singapore

**Keywords:** Huntington disease, BACHD, white matter, myelination, oligodendrocytes, germ-free, microbiota, microbiome

## Abstract

Structural and molecular myelination deficits represent early pathological features of Huntington disease (HD). Recent evidence from germ-free (GF) animals suggests a role for microbiota-gut-brain bidirectional communication in the regulation of myelination. In this study, we aimed to investigate the impact of microbiota on myelin plasticity and oligodendroglial population dynamics in the mixed-sex BACHD mouse model of HD. Ultrastructural analysis of myelin in the corpus callosum revealed alterations of myelin thickness in BACHD GF compared to specific-pathogen free (SPF) mice, whereas no differences were observed between wild-type (WT) groups. In contrast, myelin compaction was altered in all groups when compared to WT SPF animals. Levels of myelin-related proteins were generally reduced, and the number of mature oligodendrocytes was decreased in the prefrontal cortex under GF compared to SPF conditions, regardless of genotype. Minor differences in commensal bacteria at the family and genera levels were found in the gut microbiota of BACHD and WT animals housed in standard living conditions. Our findings indicate complex effects of a germ-free status on myelin-related characteristics, and highlight the adaptive properties of myelination as a result of environmental manipulation.

## INTRODUCTION

Huntington disease (HD) is a progressive autosomal-dominant neurodegenerative disorder, characterised by a triad of clinical features that comprise of progressive motor, cognitive, and psychiatric decline (Ross and Tabrizi, 2011). HD is caused by the expansion of cytosine-adenine-guanine (CAG) triplet repeats in the huntingtin (*HTT*) gene (MacDonald et al., 1993). Beyond 36 CAG repeats, the repeat length is inversely correlated with the age of motor onset (Andrew et al., 1993; Duyao et al., 1993; Snell et al., 1993). This relationship accounts for approximately 50-70% of the observed variance in clinical onset, with greater onset variance associated with lower CAG repeats, potentially due to a greater influence of genetic and environmental modifiers (Langbehn et al., 2004). It is therefore critical to investigate possible modulating environmental factors of plastic and adaptive processes contributing to HD pathology. Understanding these dynamics could contribute to the identification of approaches to modify disease onset and progression.

In addition to the well-studied neuronal degeneration in the basal ganglia, white matter (WM) atrophy and myelination defects have been identified as early features of the HD phenotype in both patients and animal models. Longitudinal studies in pre-symptomatic and early HD gene carriers have reported progressive reduction in WM volume, and indicated loss of connectivity between cortical and sub-cortical regions, which may underpin early clinical symptoms (Tabrizi et al., 2009; Tabrizi et al., 2013; Shaffer et al., 2017). White matter alterations were found in the corpus callosum (CC) and prefrontal cortex (PFC) of pre-symptomatic and symptomatic HD gene carriers (Ross and Tabrizi, 2011; Matsui et al., 2014; Matsui et al., 2015; Bourbon-Teles et al., 2017). Previous work in our laboratory has also explored the molecular and ultrastructural features of WM abnormalities in HD mouse and rat models (Garcia-Miralles et al., 2016; Teo et al., 2016). For example, diffusion tensor imaging revealed WM microstructural abnormalities prior to neuronal loss in the CC of the YAC128 mouse model and the BACHD rat model (Teo et al., 2016). In addition, thinner myelin sheaths and lower levels of myelin-related gene transcripts were seen in these animals compared to wild-type (WT) controls (Teo et al., 2016). Similar abnormalities have also been reported in the BACHD, HdhQ250 and R6/2 mouse models of HD (Wade et al., 2008; Xiang et al., 2011; Jin et al., 2015). Furthermore, transgenic mice expressing mutant HTT selectively in oligodendrocytes showed impaired myelination, with myelin thinning in the striatum, reduction of myelin basic protein gene expression, as well as progressive motor impairments, metabolic deficits and reduced survival (Huang et al., 2015).

Communication between the central nervous system, the gastrointestinal tract and gut microbiota is bidirectional and reciprocal (reviewed in Kundu et al., 2017). Notably, emerging evidence suggests the microbiota-gut-brain axis to be involved in modulation of adaptive myelination, and myelin-related characteristics, in the rodent brain (Gacias et al., 2016; Hoban et al., 2016; Lu et al., 2018). Specifically, upregulation of genes and transcripts linked to myelination have been recently described in the PFC of WT germ-free (GF) mice (Hoban et al., 2016), as well as in antibiotic treated non-obese diabetic mice (Gacias et al., 2016). These transcriptional changes were accompanied by hypermyelination of PFC axons in these animals, compared to animals raised in specific pathogen free (SPF) conditions (Hoban et al., 2016).

In the present study, we used microbiota manipulation as an environmental intervention to investigate white matter plasticity in HD. We hypothesised that the absence of microbiota differentially affects myelination in the BACHD mouse model of HD relative to wild-type controls. Our results revealed minor differences in gut bacterial populations between the two genotypes, suggesting defects in myelin plasticity observed in BACHD mice can manifest in the absence of major disturbances in gut microbiota.

## MATERIALS AND METHODS

### Experimental design

WT and BACHD transgenic mice on the FVB background were bred either under normal laboratory conditions (specific pathogen free, SPF) or in a germ-free environment (GF). GF animals were born through caesarean section in aseptic conditions, where all incoming air, water and food were sterilised. SPFs were conventional laboratory animals that possessed a naturally acquired microbiota. Both males and female mice were euthanized at 3 months of age. Mice were bred and housed in the Germ Free Animal Facility of the Singapore General Hospital Unit (SGH), Singapore. All experimental procedures were approved by, and conducted in accordance with the ethical guidelines of the animal care committee at our institution.

### Tissue extraction and processing for transmission electron microscopy (TEM)

Animals (n = 3 - 4 per group) were anesthetized with Ketamine (150mg /Kg) and Xylazine (10 mg/Kg), and transcardially perfused with phosphate buffer solution (PBS, made in-house), followed by TEM fixative solution: 2.5% paraformaldehyde (PFA, Sigma-Aldrich) and 2.5% glutaraldehyde (GlutAH, Sigma-Aldrich). Brains were placed in TEM fixative solution and stored at 4° C for 24 hours, washed in PBS and transferred in 5% sucrose (First Base) and 0.1% sodium azide (NaN_3_, Merck) at 4° C. A 1mm coronal section was made at the intersection between the CC and fornix, followed by microdissection of the anterior mid-body region of the CC (∼Bregma 1.10-0.50mm, according to the Mouse Brain Atlas, Paxinos and Franklin, 2001). Samples were post fixed in 1% osmium tetroxide (1g of osmium in 100ml PBS, pH 7.4) for 1 hour at room temperature (RT), and washed in deionised water (2×10 min). This was followed by dehydration, through an ascending ethanol series, consisting of 25%, 50%, 75%, 95%, and 100% ethanol (2×10 min, RT), and incubation in 100% acetone (2×10 min). Samples were then infiltrated with 100% acetone:resin (1:1 for 30 min at RT, and 1:6 overnight at RT), incubated with fresh resin (30 min at 40° C, and 1 hour at 45° C and 50°C, respectively), and embedded in fresh resin to polymerise (24 hours at 60°C). Following ultra-thin sectioning (90nm thickness, Diatome, Ultra45, 3mm length), samples were stained with lead citrate (Reynolds lead citrate, composed of lead nitrate and sodium citrate, 3%; for 8 minutes at RT). Samples were viewed and imaged using a Tecnai G2 Spirit Twin/ Biotwin TEM model (FEI), with the following imaging parameters: SA2550x for g-ratios, and SA11500x for periodicity.

### Ultrastructural analysis

A total of 12 images per animal were analysed using ImageJ software (version 2.0.0, National Institutes of Health). An unbiased frame was randomly selected using the rectangular tool and superimposed on each image. G-ratio of regular shaped (round) axons were obtained by calculating the ratio between the diameter of an axon and the same axon diameter plus the outer most layer of myelin surrounding that axon. Periodicity of myelin lamellae was calculated as the ratio between myelin sheath thickness and the number of major dense lines. Additionally, over 4,500 axons per group were morphologically categorised using a MATLAB (RRID:SCR_001622) custom written script.

### qRT PCR

Total RNA was extracted from mouse prefrontal cortex (n = 7 - 8 animals per group) using trizol (Life Technologies) and subsequently an RNeasy plus mini kit (Qiagen) according to the manufacturer’s insructions. First-strand cDNA synthesis was performed using the SuperScript First Strand Synthesis System (Invitrogen). Gene expression was analysed by qPCR using Sybr Select Universal Master Mix (Invitrogen) on a StepOnePlus to obtain Ct values. Each sample of every group was run in triplicate and the expression levels were calculated as the average of these replicates relative to β-actin expression. The mouse-specific primers for beta-actin, myelin basic protein (*Mbp*), as 2’,3’-cyclic-nucleotide 3’- phosphodiesterase (*Cnp*), myelin-oligodendrocyte glycoprotein (*Mog*), and Ermin (*Ermn*) are summarized in Table 1.

**Table 1:**
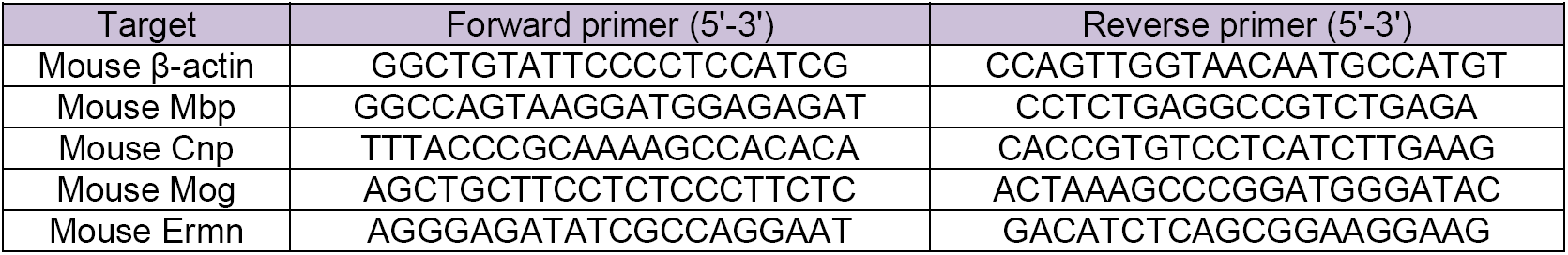
Primers used for cDNA analyses

### Western blot processing and quantification

The PFC (n = 4 animals per group) was microdissected, snap-frozen in liquid nitrogen and stored at −80°C until further use. The tissue was homogenized in RIPA buffer (Sigma-Aldrich), with PMSF (1mM, Sigma-Aldrich), Z-VAD (5µm, Promega), NaVan (1mM, Sigma-Aldrich), and Complete Protease Inhibitor Cocktail tablets (Roche). Protein concentration was measured using the Bradford assay (BioRad). The samples were denatured (70°C, 10 min) in 4x NuPage sample buffer and 10x NuPage Reducing agent (Thermo Fisher). A 10 well, 12% Bis-Tris protein gel (Novex) was used to separate 20µg of protein (100V, 3 hours), followed by transfer to nitrocellulose membrane (120V, 1.5 hours, RT). The samples were probed for myelin basic protein (MBP) (1:5000; Millipore Cat# MAB386, RRID:AB_94975), proteolipid protein (PLP) (1:1000; Abcam Cat# ab28486, RRID:AB_776593) and Ermin (Ermn) (1:2000; Merck Cat# MABN323). Endogenous controls used were rabbit anti-Calnexin (1:5000, Sigma-Aldrich Cat# C4731, RRID:AB_476845) and mouse anti-β-actin (1:5000, Sigma-Aldrich Cat# A5316, RRID:AB_476743). Secondary antibodies (1:10,000) included: Alexa-Fluor goat anti-rabbit 680 (Thermo Fisher Scientific Cat# A27042, RRID:AB_2536103), Alexa-Fluor goat anti-mouse 800 (Thermo Fisher Scientific Cat# A32730, RRID:AB_2633279), and Alexa-Fluor goat anti-rat 800 (Thermo Fisher Scientific Cat# SA5-10024, RRID:AB_2556604). Membranes were imaged using the LiCor Imaging System and Odyssey V3.0 software (LiCor), and blots were quantified using Image Studio Lite.

### Tissue processing for immunostaining

Mice were anaesthetised with intraperitoneal injections of ketamine (150 mg/kg)/xylazine (10 mg/kg) mixture, and brains were extracted and post-fixed in 4% PFA solution (4g in 100ml 1xPBS) for 24 hours, and subsequently submerged in 30% sucrose & 0.1% NaN3 in 1xPBS. Brains were cryopreserved in dry ice and isopentane (2-methylbutane), embedded onto chucks using optimum cutting temperature (OCT) compound (Sakura, Torrance, CA, USA), and coronal slices were sectioned at 25µm using a cryostat (Microm HM 525, Thermo Fisher Scientific, Waltham, Massachusetts, USA). Collection began when the forceps minor of the corpus callosum became visible during sectioning. Following sectioning, slices were stored in 1xPBS and 0.1% NaN3 solution at 4°C.

### Immunohistochemistry

A series of coronal sections spanning the corpus callosum were stained with oligodendroglial lineage marker (Olig2; 1:300; Millipore Cat# AB9610, RRID:AB_570666), and Glutathione S-transferase (GST-pi; 1:1000; MBL International Cat# 311, RRID:AB_591791) in 5% normal goat serum (NGS) 0.2% Triton X-100 in PBS (overnight, 4°C). This was followed by incubation in biotinylated secondary antibody (1% NGS, 0.2% Triton, 1:200 ABC Vectastain Kit solution (rabbit kit, Vector Laboratories Cat# PK-6100, RRID:AB_2336819); 1.5 hours, RT), three washes in 1xPBS, and incubation in Vectastain Elite ABC reagent (2 hours, RT). Sections were stained using 3,3′-diaminobenzidine DAB (ImmPACT DAB Peroxidase Substrate, Vector Laboratories Cat# SK-4105, RRID:AB_2336520), mounted on slides and cover slipped with DPX mounting media (Sigma Cat# 44581).

### Immunofluorescence

Sections were treated for antigen retrieval in hydrochloric acid (HCl) based solution (stock prepared from 37% HCl solution): 1N HCl (10 min, 4°C), 2N HCl (10 min, RT), 2N HCl (20 min, 37°C), and 0.1M borate buffer solution (pH 9; 10 min, RT). A standard staining protocol was then followed where sections were sequentially washed (1xPBS; 3×10 min), incubated in blocking solution (5% normal donkey serum - NDS, Sigma-Aldrich; 0.2% Triton; in PBS; RT, 120 min), incubated in primary antibody solution (1% NDS, 0.2% Triton, in PBS; overnight; RT), and stained with goat anti-platelet derived growth factor receptor alpha (PDGFRα; 1:500; R and D Systems Cat# AF1062, RRID:AB_2236897), washed (1xPBS, 3×10 min), and incubated with donkey anti-goat Alexa-488 (1:500 in PBS; Thermo Fisher Scientific Cat# A-11055, RRID:AB_2534102) with 0.2% Triton in PBS (2 hours; RT). Sections were mounted on slides and cover slipped with Fluoromount(tm) mounting medium (Merck KGaA).

### Stereology

Stereo Investigator software (MBF Bioscience) with optical fractionator connected to an AxioImager M2 microscope (Carl Zeiss AG) and AxioCam MRc Digital CCD camera (Carl Zeiss AG) was used for imaging and cell counting. The coronal navigation of the mouse brain atlas (Paxinos and Franklin, 2001) was used as reference to draw contours for the regions of interest: PFC (Bregma 1.94 to 1.18mm) and CC (Bregma 1.18 to −1.94mm). Section thickness was set to 25µm and section evaluation interval was set to 12 (every 12th section was selected for counting). A total of 9-12 animals per condition, and 8-12 sections per animal were used. For GST-pi and Olig2, the counting frame size was set to 100×100µm for the PFC and 50×50µm for the CC; while for PDGFRα the counting frame size was set to 250×250µm for both regions. An estimated grid size, with a total of 15 sampling sites, was selected for counting cells for both regions. For all measurements a 10µm dissector height and 5µm guard were used. The precision of the estimates was determined by using the Gundersen and Schmitz-Hof coefficient of errors for each section.

### Faecal DNA extraction, and bacterial 16S rRNA sequencing and processing

Faecal samples were collected from 3 months of age (n = 10 per group) and 6 months of age (n = 5 - 6 per group) mixed sex BACHD SPF and WT SPF mice. Extraction of faecal genomic DNA was performed through a combination of chemical and mechanical lysis methods, using FastPrep Instrument (MP Biomedicals) and QIAamp Fast DNA Stool Mini Kit (Qiagen Cat# 51604). From total DNA, the V3-V6 region of the prokaryotic 16S rRNA gene was amplified as previously described (Ong et al., 2013; The et al., 2017). Amplicons were sheared using the Covaris LE220 sonicator (Covaris Inc.), and sequencing libraries were built using GeneRead DNA Library I Core Kit (Qiagen Cat# 180434). DNA libraries were multiplexed by 96 indices, pooled, and sequenced on the Illumina HiSeq 2500 using paired-end (2×76bp) sequencing. Sequencing reads were de-multiplexed (Illumina bcl2fastq 2.17.1.14 software) and filtered (PF=0) before conversion to FASTQ format. Sequence clean-up was performed by removing trailing bases (quality score ≤ 2; reads ≤ 60bp; Miller et al., 2011). Full length 16S rRNA reconstructions were obtained from the short sequencing reads (Expectation Maximisation Iterative Reconstruction of Genes from the Environment (EMIRGE) amplicon algorithm; Miller et al., 2013). Reconstructed sequences that have at least 99% sequence similarity were collapsed into operational taxonomic units (OTUs), and Graphmap was used to map these OTUs to the Greengenes global rRNA database (greengenes/13_5/99_otus.fasta; DeSantis et al., 2007). The analysis is based on the method described in Ong et al. (2013) with updated tools (https://github.com/CSB5/GERMS_16S_pipeline). OTUs were called at various levels of identity (species, genus, family) dependent on sequence similarity to the 16S rRNA sequence entries for bacteria in the Greengenes (RRID:SCR_002830) and SILVA (RRID:SCR_006423) databases (Yarza et al., 2014). Hits below predefined percent identity (97% at the species level, 94.5% at the genus level, 86.5% at the family level and 75% at the phylum level) were excluded from classification. The relative abundance of OTUs was obtained, and transformed to relative abundances at the phylum, family, genus and species level.

### Microbiota populations analysis

To assess differences in microbiota populations between the BACHD and WT groups, distance-based redundancy analysis (dbRDA), based on Bray-Curtis distances with the capscale function of vegan package in R (R Project for Statistical Computing, RRID:SCR_001905) was used. Permutational Multivariate Analysis of Variance (PERMANOVA) was applied based on 999 permutations to determine if microbial community composition differences were significant between the two groups. Relative abundance differences between the two groups were statistically assessed by Mann-Whitney U-test. At individual bacterial family and genera level no significant differences were identified following corrections for multiple comparisons, thus, nominal p-values were used to identify potential differences and trends. Mean ± SD values were used to describe the results, while mean ± SEM values were used for creating the bar graphs.

### Statistical analysis

Two-way ANOVA with Sidak post hoc test of correction for multiple comparisons was used for the 2×2 study design (genotype x microbiota presence) to observe whether differences between the four groups were statistically significant, as appropriate. Cumulative frequencies of g-ratios were analysed using the unpaired two-tailed Kolmogorov-Smirnov test. For western-blot analysis, pairwise comparisons were generated using Student’s two-tailed t-tests. For all statistical tests, p-values < 0.05 were considered statistically significant. P-values and n-values were indicated in the associated legends for each figure, while mean ± SEM values were stated in the results section. Bar graphs were created based on mean ± SEM, unless otherwise stated. All bar graphs and statistical results were obtained using Prism 7, GraphPad Prism (RRID:SCR_002798).

## RESULTS

### Differential effects of microbiota manipulation on myelin and axonal characteristics in the CC of BACHD and WT animals

To examine the effects of a germ-free environment on myelination in BACHD and WT animals, we elected to investigate ultrastructural myelin and axonal characteristics in the CC using transmission electron microscopy (TEM, Fig. 1A). Myelin thickness of regular shaped axons was quantified using g-ratios, defined as the ratio between axonal diameter to axonal diameter plus myelin lamellae (Friede and Miyagishi, 1972). G-ratios were first assessed as a function of axon diameter in order to examine variability of myelin thickness at different axonal calibre ranges, and to identify any abnormalities relative to control (Fig. 1B). Comparison of WT groups showed fairly similar g-ratios over most axonal diameter ranges for SPF and GF animals (Fig. 1B, top), whereas BACHD GF animals had lower g-ratios compared to BACHD SPF (Fig. 1B, bottom), particularly at small to mid-range calibre axons (< ∼1000nm). In addition, cumulative frequency analysis of g-ratios of both SPF and GF WT groups exhibited a close overlap (U = 539,054; p > 0.05, two-tailed Mann-Whitney test; Fig. 1C, top), indicating similarity between the two groups, while an evident shift towards smaller values was observed for BACHD GF compared to BACHD SPF (U = 558,086; p < 0.0001, two-tailed Mann-Whitney test; Fig. 1C, bottom). To further examine changes in myelin thickness as a function of axon calibre, g-ratio values were subsequently classified, averaged, and compared according to three axon calibre ranges: small (< 500nm), mid-range (500 ≤ diameter < 1000nm), and large axons (≥ 1000nm) (Fig. 1D). WT GF animals showed no differences in mean g-ratio compared to WT SPF animals in mid-range and large axons (Fig. 1D; mid-range WT GF: 0.847±0.002, WT SPF: 0.846±0.001; large WT GF: 0.875±0.005, WT SPF: 0.895±0.002), although a significant increase in g-ratios was observed at small axonal diameters (Fig. 1D; WT GF: 0.805±0.002; WT SPF: 0.782±0.002; microbiota presence effect, F(1, 1869) = 6.96, p = 0.0084, two-way ANOVA; t(1869) = 4.94, p < 0.001, Sidak *post hoc* test). On the other hand, BACHD GF animals had smaller g-ratios when compared to BACHD SPF (Fig. 1D), which was most evident at both small (GF: 0.752±0.003; SPF: 0.789±0.002; t(1869) = 10.13, p < 0.001, Sidak *post hoc* test) and mid-range axonal diameters (GF: 0.824±0.002; SPF: 0.841±0.001; microbiota presence effect, F(1, 2601) = 13.36, p = 0.0003, two-way ANOVA; t(2601) = 5.61, p < 0.001, Sidak *post hoc* test), and to a lesser degree at large diameter axons, although the same trend was still observed (GF: 0.865±0.007; SPF: 0.878±0.003; p > 0.05, Sidak *post hoc* test). This classification of g-ratios based on axonal calibre indicates thinner myelin for small axons in WT animals, and thicker myelin at all axonal calibre ranges for BACHD animals placed in a germ-free environment. Finally, when averaged over all axonal diameters (Fig. 1E), a significant interaction between genotype and microbiota presence was observed (two-way ANOVA: interaction effect, F(1, 4374) = 121.7, p < 0.0001; microbiota presence effect, F(1, 4374) = 72.3; p < 0.0001; genotype effect, F(1, 4374) = 175.8; p < 0.0001). G-ratios of both SPF and GF WT groups were comparable (p > 0.05, Sidak *post hoc* test), while the mean g-ratio of BACHD GF animals (0.786±0.002) was significantly reduced compared to both WT GF (0.834±0.001; t(4374) = 15.82, p < 0.0001, Sidak *post hoc* test), and BACHD SPF (0.824±0.001; t(4374) = 14.05, p < 0.0001, Sidak *post hoc* test) (Fig. 1E). This detailed analysis of g-ratio changes as a function of axonal diameter suggests that the alterations in callosal myelination was more pronounced in BACHD mice compared with WT mice bred in GF housing conditions (summarised in Table 2).

**Table 2.**
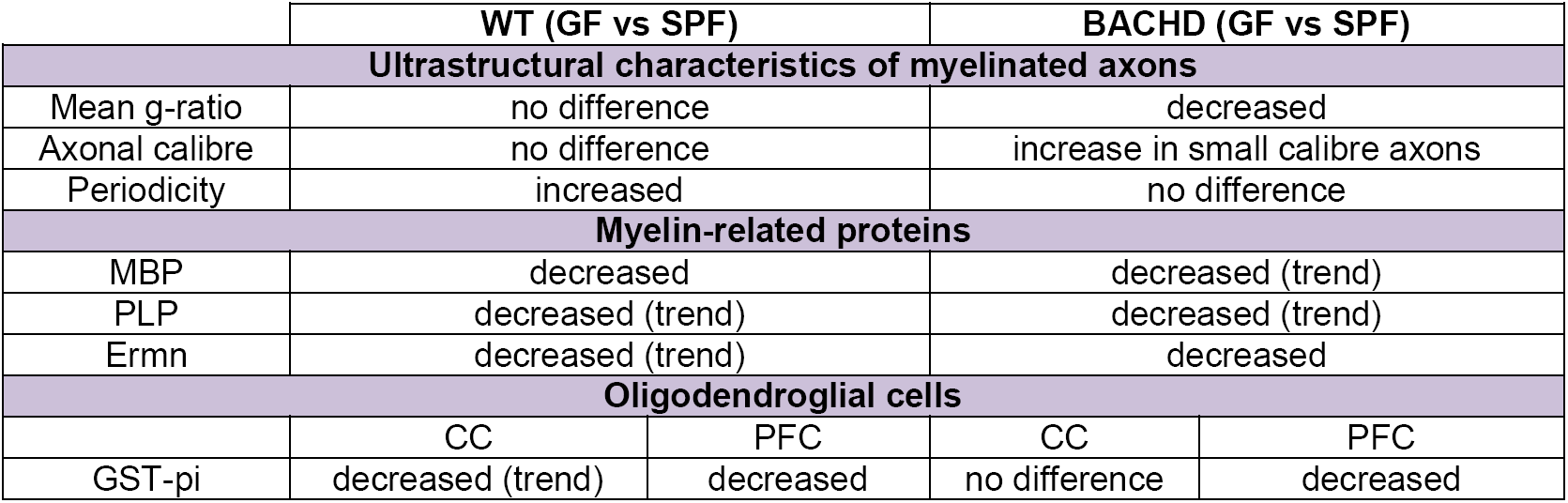
Summary of findings

**Fig 1.**
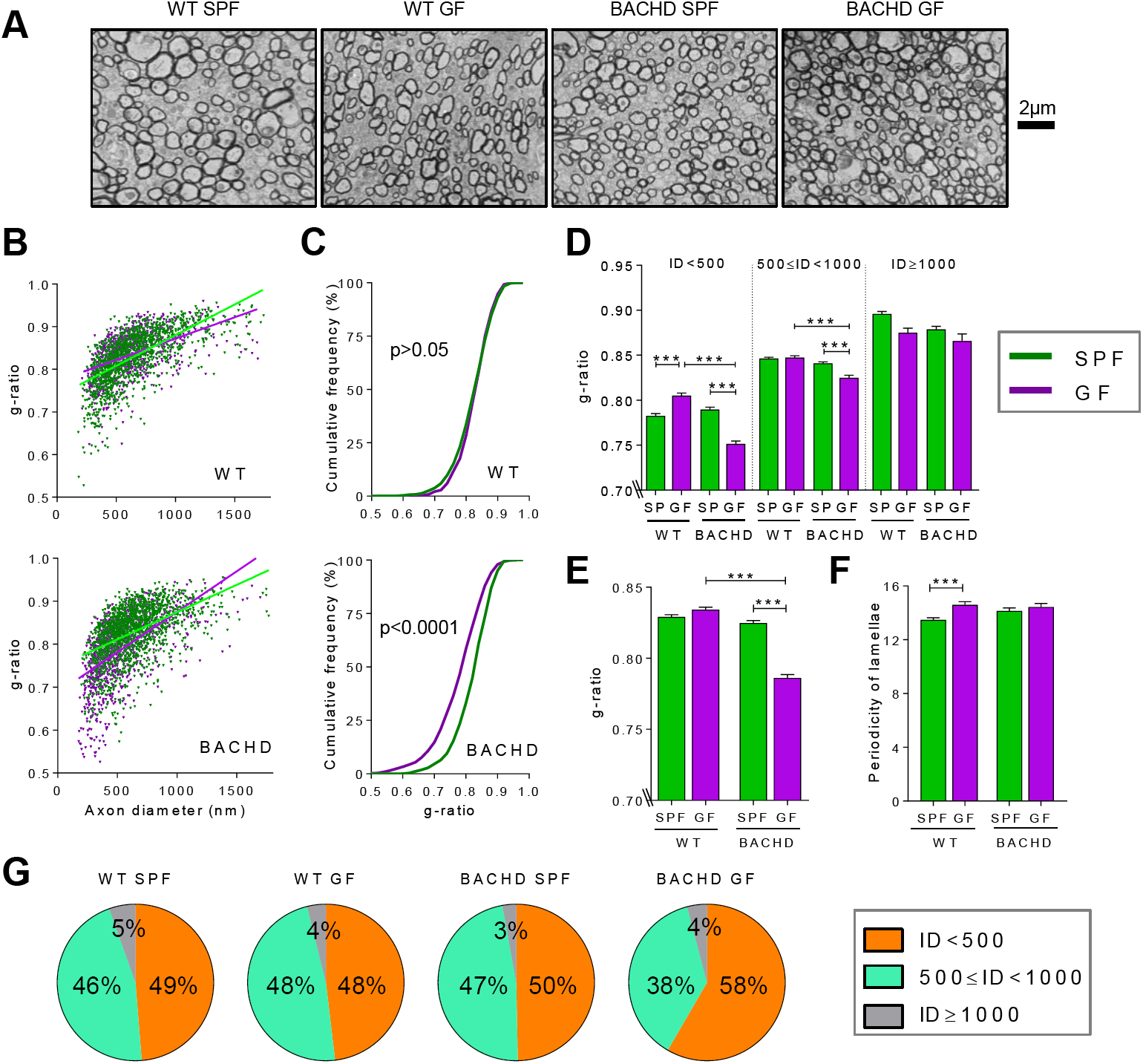
Differential effects of a germ-free status on myelination and axonal characteristics in the corpus callosum (CC) of BACHD and WT mice. (A) Resentative micrographs of axons from the anterior-mid region of the CC for each group.scale bar 2 μm. (B) Scatter plot of g-ratios as a function of axons diameter. (C) Cumulative frequency plot of g-ratio for WT (top), and BACHD (bottom) (D) G-ratios classified according to axonal diameter (small, ID < 500nm; mid-range, 500 ≤ ID < 1000nm; large, ID ≥ 1000nm). (E) Mean g-ratio at all axonal diameters. Axons, n = 775-1440 per condition (B-E). (F) Periodicity, calculated as the ratio between myelin thickness and number of major dense lines. Axons, n = 174-358 per condition. (G) Pie chart representation of small, mid- and large size myelinated axons. Axons, n = 4,890-6,878 per condition. Bar graphs represent mean ± SEM. For all graphs, n = 3-4 mice per condition were used. Two-way ANOVA, with Sidak’s multiple comparisons test (D, E) and two-tailed Mann-Whitney test, with Bonferroni corrections (F). *p<0.05; ***p < 0.001; ****p < 0.0001. Axon diameter and periodicity shown in nanometres (nm). Abbreviations: GF, germ-free; SPF, specific-pathogen free.

On average myelin sheaths appeared to be thicker in BACHD animals housed in a GF environment, indicating that removing microbiota may contribute to hypermyelination. However, myelin ultrastructure differences were further identified by investigating the compaction of myelin membranes (Fig. 1F). By EM, compact myelin is viewed as a lamellar structure of alternating dark (major dense) and light (intraperiod) lines. Periodicity was defined as the ratio between myelin sheath thickness and the number of major dense lines. Increased periodicity of myelin lamellae, suggestive of reduced compaction, was observed in the mean of all groups (BACHD SPF: 14.14±0.23 nm; BACHD GF: 14.44±0.26 nm; WT GF: 14.59±0.24 nm) compared to the WT SPF group (13.47±0.16 nm). This was statistically significant when comparing WT GF and WT SPF mice (U = 31,862, p = 0.0006, two-tailed Mann-Whitney test, with Bonferroni correction for multiple comparisons), and a trend was observed between BACHD SPF and WT SPF mice (U = 35,177, p = 0.03, two-tailed Mann-Whitney test, Bonferroni correction requiring p ≤ 0.0125) (Fig. 1F). Therefore, increased periodicity suggests decompaction of myelin lamellae as both an effect of a GF status, but also as a possible effect of the *HTT* mutation. Additionally, the apparent increase in myelin sheath thickness observed in small calibre axons of BACHD GF animals, as implied by the decrease in g-ratios, may be partly due to decompaction.

Over 4,500 axons per group were morphologically categorised to further characterise the effects of germ-free manipulation on myelinated fibres in the CC of BACHD and WT animals (Fig. 1G). Following classification of axons based on diameter size, BACHD GF animals were found to possess an increased proportion (58%) of small diameter axons (<500nm) compared to all other groups (48-50%) (Fig. 1G). In turn, the proportion of mid-range diameter axons in BACHD GF (38%) was found to be markedly reduced relative to other groups (46-48%) (Fig. 1G). No differences in the proportion of large diameter axons were observed between the groups (3-5%). These findings suggest an increase in representation of small axons that were myelinated in the BACHD GF animals compared to the other groups.

### Decreased levels of myelin-related proteins in the PFC of mice raised in a germ-free environment

In order to understand whether differences in myelin and axonal characteristics, observed at ultrastructural level correlate with changes at gene or protein level, candidate myelin-related genes and associated proteins were evaluated. Since myelination in the PFC has been previously speculated to be particularly plastic and susceptible to microbiota manipulation, we chose to investigate this region. Four myelin specific genes, involved in maintaining stability, integrity and function of the myelin sheath, were quantified using qRT-PCR analysis (Fig. 2A). No significant differences were observed in levels of *Mbp* between GF and SPF conditions in either of the two genotypes (two-way ANOVA with Sidak *post hoc* test; p > 0.05; Fig. 2A). Other myelin-related genes such as *Cnp, Mog*, and *Ermn* showed broadly similar levels of expression across all four groups. Quantification by western blot of myelin specific proteins (MBP, PLP and Ermn) revealed significant, or trends towards, decreased levels in GF groups compared with their respective SPF groups for both genotypes (MBP, t(6) = 3.03, p = 0.023; Ermn, t(6) = 3.67, p = 0.01; Student’s two-tailed t-test; Fig. 2B). MBP protein band 14kDa was quantified and illustrated here, while the other three isoforms (18kDa, 21.5kDa and 24kDa) reflected changes in the same direction. These results suggest that decreased levels of compact myelin-related proteins, such as MBP and PLP in WT GF animals, may contribute to the increased periodicity of myelin sheaths as a result of the germ-free environment (summarised in Table 2).

**Fig 2.**
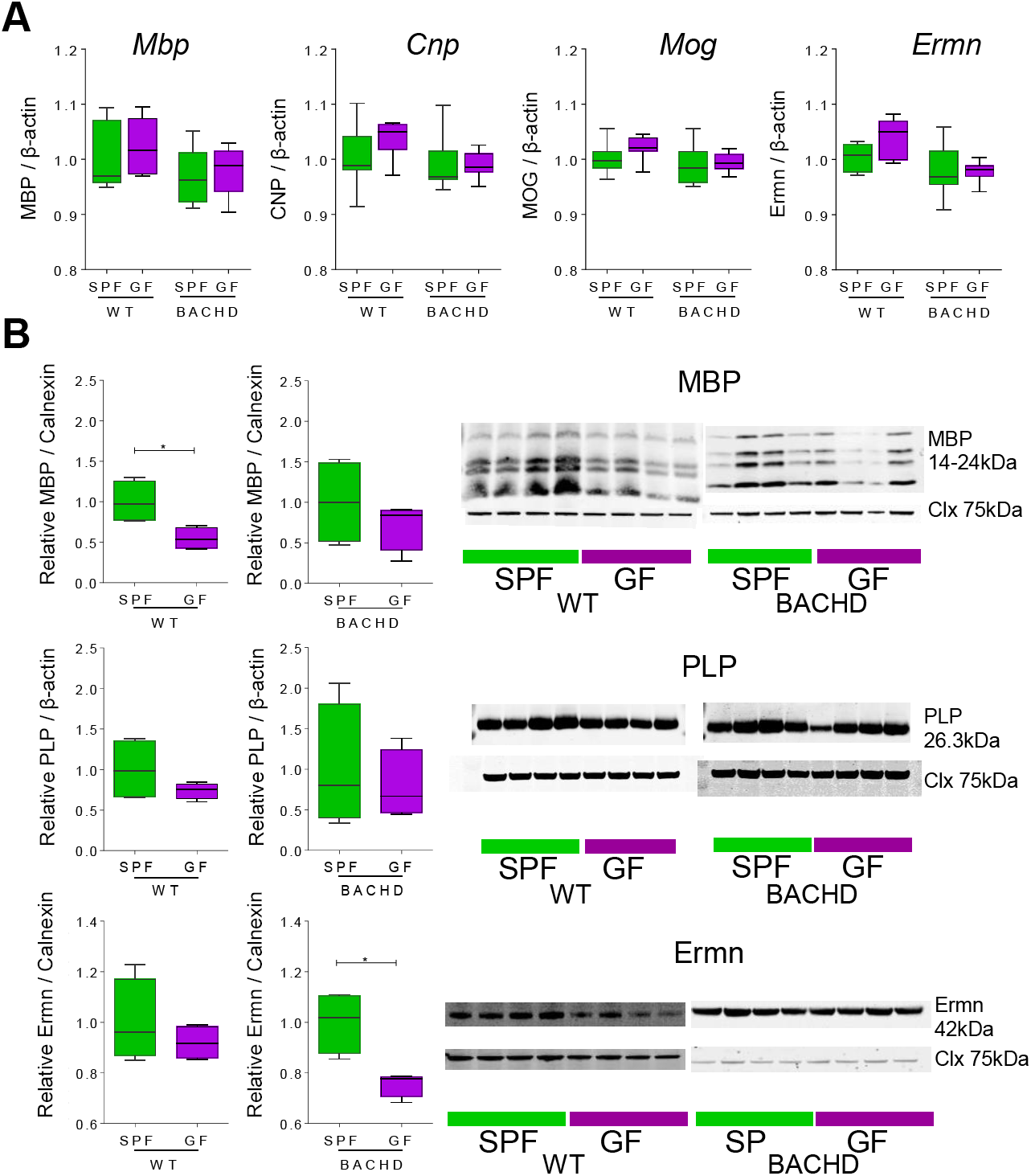
Myelin transcripts and protein levels in the prefrontal cortex (PFC) of BACHD and WT mice. (A) Myelin transcripts levels: *Mbp, Cnp, Mog*, and *Ermn* (from left to right). Values of qPCR results were normalised to β-actin mRNA levels and are represented as fold change relative to SPF values. Bar graphs represent mean ± SEM (n = 7-8 mice per group). Two-way ANOVA, with Sidak’s multiple comparisons test. (B) Quantification of relative protein levels and their representative western blots of PFC homogenates immunostained with MBP and Calnexin (top), PLP and β-actin (middle), and Ermn and Calnexin (bottom). Quantification of each protein of interest was normalised to Calnexin or β-actin, and expressed relative to SPF control for each blot. Bar graphs represent mean ± SEM (n = 4 mice per group). Student’s unpaired two-tailed t-test, 95% confidence intervals; *p < 0.05. Abbreviations: Clx, calnexin; CNP, 2’,3’-cyclic-nucleotide 3’-phosphodiesterase; Ermn, Ermin; GF, germ-free; MBP, myelin basic protein; MOG, myelin-oligodendrocyte glycoprotein; PLP, proteolipid protein; SPF, specific-pathogen free.

### Reduction of mature oligodendroglial population as a result of germ-free status

Oligodendroglial populations were quantified at different stages of development in both the CC and PFC of BACHD and WT mice under SPF or GF conditions. The oligodendrocyte transcription factor, Olig2, is expressed when cells become committed to the oligodendroglial lineage, (Lu et al., 2002), and is downregulated with oligodendrocytes maturation (Ligon et al., 2006). Oligodendrocyte precursor cells (OPCs) express platelet derived growth factor receptor alpha (PDGFRα; Pringle et al., 1992), whereas the cytoplasm of mature myelinating oligodendrocytes contains, among others, Glutathione S-transferase of class pi (GST-pi; Cammer et al., 1989). There were no significant differences between the groups in PDGFRα or Olig2 positive cells numbers in the CC region (Fig. 3A, left and middle panels). However, there was a trend towards a decreased number of GST-pi positive cells in the CC region of WT GF animals compared to SPF controls (WT GF (21,027±2,570) vs WT SPF (31,360±5,445); two-way ANOVA with Sidak *post hoc* test; p > 0.05; Fig. 3A, right panel). Similarly to the CC, there were no significant differences in either PDGFRα or Olig2 numbers in the PFC region between the four groups (Fig. 3B, left and middle panels). Interestingly, there was a significant effect of the GF status on GST-pi positive cells in the PFC (F(1,35) = 24.46, p < 0.0001, two-way ANOVA; Fig. 3B, right panel), with this population decreased for both GF groups compared to SPF controls (WT GF (3,318±412) vs WT SPF (8574±1291); t(35) = 4.22, p = 0.0003, Sidak *post hoc* test; and BACHD GF (4,430±352) vs BACHD SPF (8,180±720); t(35) = 2.82, p = 0.015, Sidak *post hoc* test). Additionally, Olig2 positive cells from both the CC (Fig. 3A, middle panel) and PFC (Fig. 3B, middle panel) regions exhibited a trend towards reduced numbers in WT GF animals (CC: 23,263±2,261; PFC: 8,228±805.9) compared to SPF controls (CC: 38,460±4,630; PFC: 12,937±1,633; two-way ANOVA with Sidak *post hoc* test; p > 0.05). Collectively, these results suggest a reduction of mature oligodendrocytes as a result of GF status, as reflected by the decreased number of GST-pi positive cells in the PFC and CC regions.

**Fig 3.**
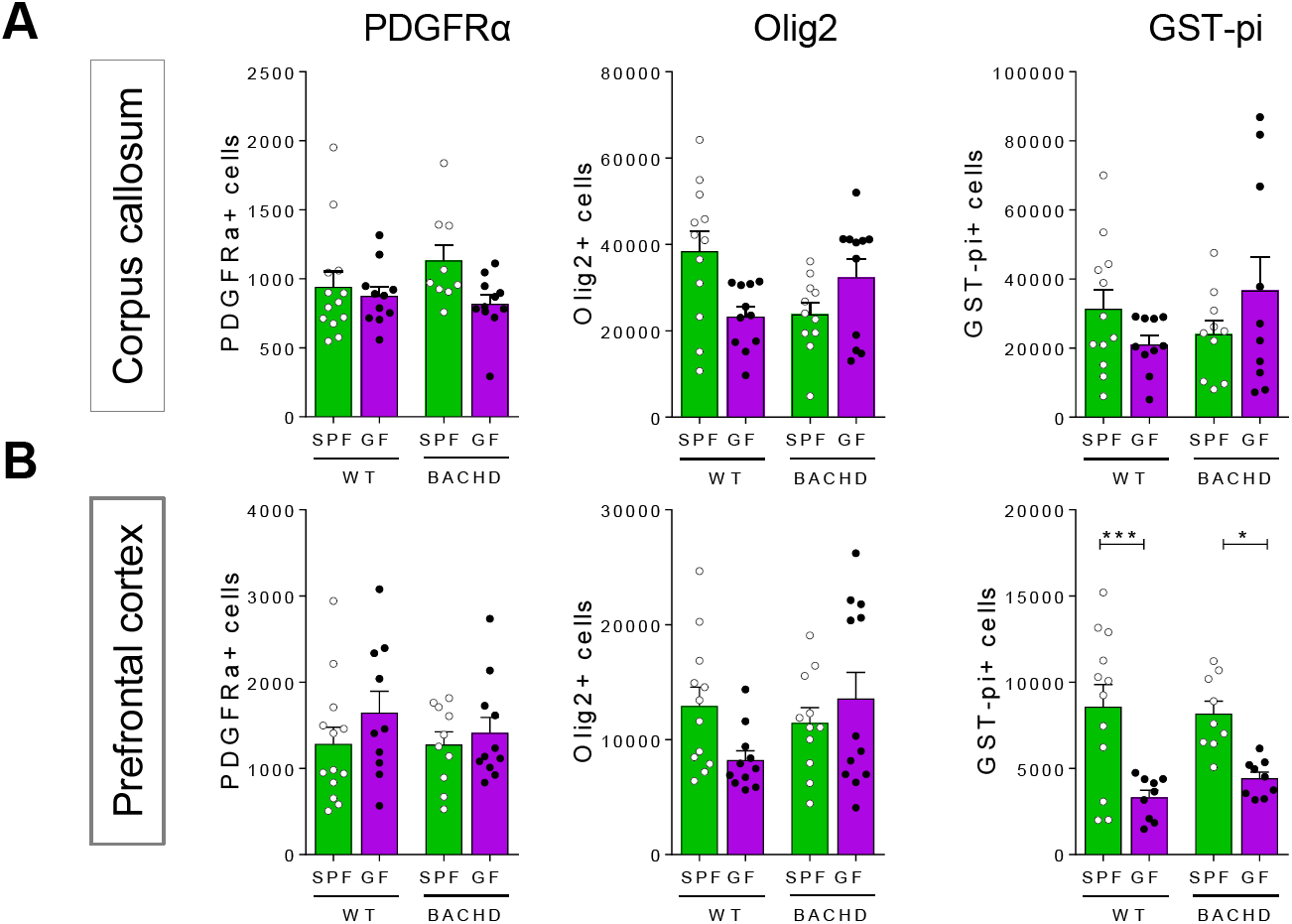
Quantification of oligodendroglia sub-populations in the CC and PFC of BACHD and WT mice. (A) Bar graphs showing stereological estimates (per mm^2^) of PDGFRα+ cells, Olig2+ cells, and GST-pi+ cells in the CC, (B) and PFC. Bar graphs represent mean ± SEM. For all graphs, n = 9-12 animals per condition, and n = 8-12 sections per animal were analysed. Two-way ANOVA, with Sidak’s multiple comparisons test. *p < 0.05; ***p < 0.001. Abbreviations: GF, germ-free; GST-pi, Glutathione S-transferase pi; Olig2, oligodendrocyte transcription factor; PDGFRα, platelet derived growth factor receptor alpha; SPF, specific-pathogen free.

### Characterisation of microbiota in BACHD and WT mice housed under SPF conditions

To identify differences between the gut microbiota community of BACHD and WT animals housed under SPF conditions, we conducted 16S rRNA gene sequencing on DNA isolated from faecal samples collected at 3 and 6 months of age (Fig. 4). The Shannon diversity index, which considers species richness and the evenness of abundance, showed a small non-significant decrease in diversity composition in BACHD animals compared to WT (not shown) at 6 months of age. Taxonomic classification revealed a profile comprising of Bacteroidetes, Firmicutes, Tenericutes, Proteobacteria, TM7 (Fig. 4A), and other rare bacterial phyla, such as Cyanobacteria, Actinobacteria and Deferribacteres (not shown). When comparing 3 months of age BACHD to WT animals (Fig. 4A, top pie charts), relative abundance at phylum level revealed a decrease in Bacteroidetes (BACHD 72.8±6% vs WT 77.5±7%), and increase in Firmicutes (25.8±6% vs 20.8±6%); whereas, at 6 months of age (Fig. 4A, bottom pie charts) the opposite effect was observed, with increased Bacteroidetes (BACHD 75.6±7% vs WT 72.5±6%), and decreased Firmicutes levels (23.1±7% vs 25.4±6%). Furthermore, when commensal bacteria was classified and compared at family and genera levels, only minor differences were identified between BACHD and WT conventional mice (Fig. 4B). At the bacterial family level (Fig. 4C), 3 month old BACHD mice showed decreased *Bacteroidaceae* (phylum Bacteroidetes; BACHD (1.05±0.48) vs WT (1.59±0.58); Mann-Whitney U-test; nominal p-value = 0.0185) and *Anaeroplasmataceae* (phylum Tenericutes; BACHD (0.02±0.02) vs WT (0.07±0.07); Mann-Whitney U-test; nominal p-value = 0.028), whereas at 6 months of age BACHD mice showed decreased *Mogibacteriaceae* (phylum Firmicutes; BACHD (0.01±0.004) vs WT (0.02±0.01); Mann-Whitney U-test; nominal p-value = 0.017). Further minor differences were identified at the genera level (Fig. 4D), where BACHD mice showed lower abundance of *Prevotella* (phylum Bacteroidetes; 3 months: BACHD (3.83±1.46) vs WT (5.91±1.76); Mann-Whitney U-test; nominal p-value = 0.018; 6 months: BACHD (4.37±2.19) vs WT (7.7±2.39); Mann-Whitney U-test; nominal p-value = 0.082) and *Bacteroides* (phylum Bacteroidetes; 3 months: BACHD (1.2±0.77) vs WT (2.51±1.01); Mann-Whitney U-test; nominal p-value = 0.003; 6 months: BACHD (1.62±0.6) vs WT (2.22±1.32); Mann-Whitney U-test; nominal p-value > 0.05) at 3 months of age, and a similar trend at 6 months of age. Higher abundance of *Oscillospira* (phylum Firmicutes; BACHD (1.28±0.51) vs WT (0.87±0.26); Mann-Whitney U-test; nominal p-value = 0.035) and *Adlercreutzia* (phylum Actinobacteria; BACHD (0.01±0.008) vs WT (0.007±0.003); Mann-Whitney U-test; nominal p-value = 0.043) was observed only at 3 months of age (Fig. 4D). Interestingly, these results suggest fairly similar microbiota profiling between BACHD and WT conventional animals housed in the same home cages.

**Fig 4.**
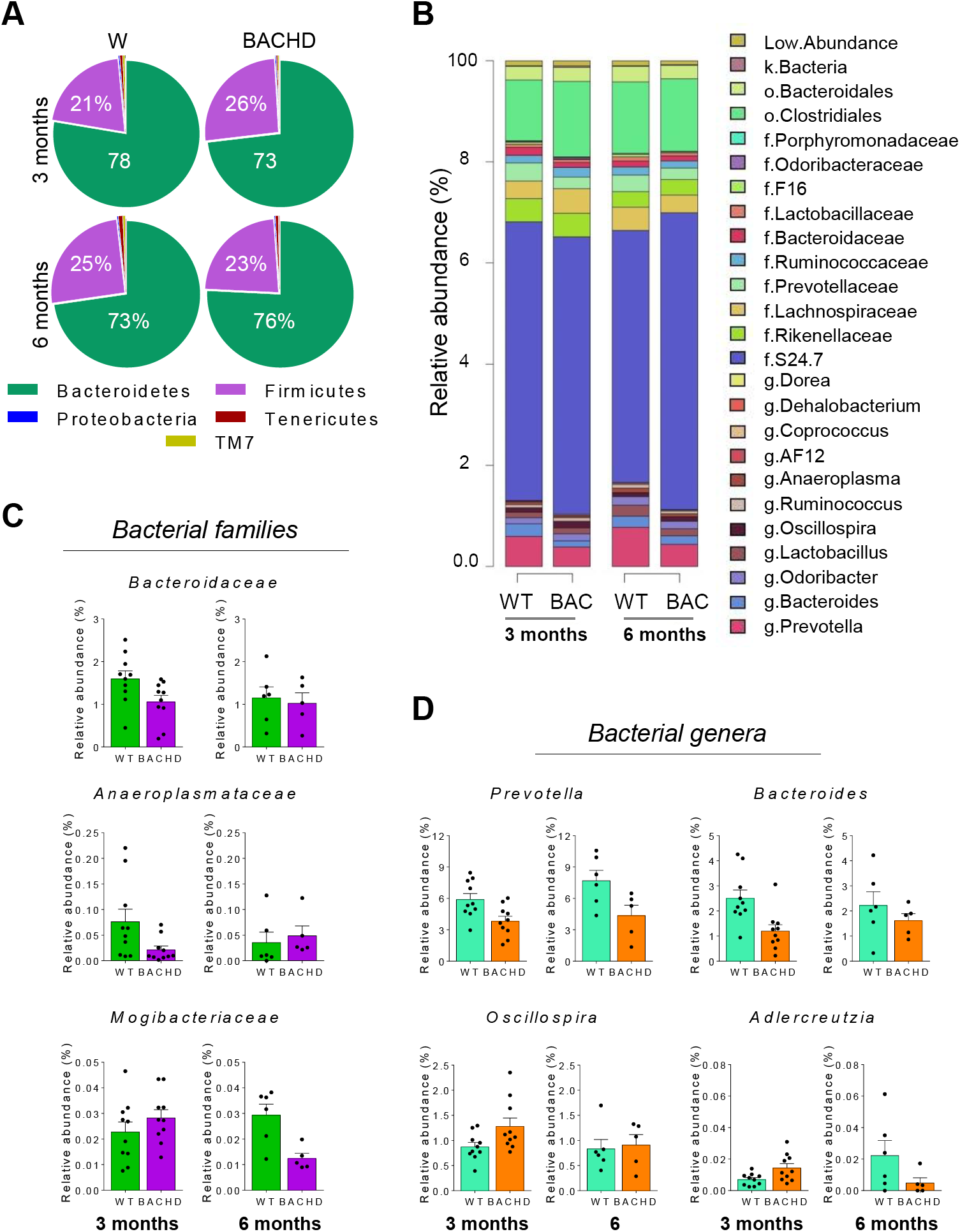
Microbiome profiling of BACHD SPF and WT SPF mice. (A) Mean relative abundance of gut microbial taxa at phylum level in BACHD and WT SPF mice, aged 3 and 6 months. (B) Relative abundance of gut microbiota at genus level. (C) Bacterial families and (D) genera differentially expressed between BACHD mice and WT control mice, either at 3 or 6 months of age (using nominal p > 0.05). Bar graphs represent mean ± SEM; n = 10 mice per group, aged 3 months; n = 5-6 mice per group, aged 6 months. Bar plots are colour coded (family: green / purple; genera: turquoise / orange). Relative abundance shown as individual data points.

## DISCUSSION

We investigated the effects of microbiota manipulation on myelination and oligodendroglial population in the BACHD mouse model of HD and WT controls. Ultrastructural analyses indicated that a GF environment had a greater effect on the extent of myelination and calibre of callosal axons in BACHD animals compared to WT controls, while possible decompaction of myelin lamellae was observed in both genotypes. Alterations of myelin-related proteins and mature oligodendrocytes as a result of GF status were observed in both genotypes. Characterisation of microbiota population in SPF animals revealed minor shifts in commensal bacteria of BACHD compared to WT controls, suggesting that mutant *HTT* in BACHD mice does not cause any notable disturbances in gut microbiota. These findings imply that defects in myelin plasticity observed in BACHD mice are not likely to be driven by disturbances in gut microbiota.

BACHD mice raised under GF conditions exhibited an 8-10% increase in the number of small calibre myelinated callosal axons, reflected by the decrease in axonal diameter size. These axons, and mid-range calibre axons, representing the majority of all axons in our analysis, were associated with lower g-ratios, suggestive of thicker myelin. However, investigation of myelin membranes indicated the expansion observed may be partly due to lamellae decompaction rather than an increase in the number of lamellae. A trend towards increased periodicity, suggestive of decompaction, was also identified in BACHD SPF compared to WT SPF animals; therefore this effect could also be caused by the *HTT* mutation in BACHD animals. Decompaction of lamellae in BACHD GF animals is supported by our observation of a trend towards lower levels of cortical MBP and PLP, which are among the most abundant specific myelin-related proteins in the CNS and play a key role in myelin compaction (Klugmann et al., 1997; Boggs, 2006; Min et al., 2009). Additionally, oligodendrocyte-specific cytoskeletal protein, Ermn, was also decreased in BACHD animals raised in a GF environment, and has been implicated to play a role in late wrapping and the compaction phase of myelination (Brockschnieder et al., 2006).

Myelin alterations in WT animals in response to a GF status, in contrast, were somewhat different to those identified in BACHD animals. There were major changes in overall axonal size, although myelin thickness was decreased at small calibre axons, with decompaction of lamellae across all axonal calibres. These findings differ from a previous study that found WT GF male mice to show hypermyelination over all axonal diameters in the PFC region (Hoban et al., 2016). This disparity may be explained by differences in the brain regions investigated (anterior mid-body CC here, and PFC in the aforementioned study). It is also possible that the PFC might be particularly susceptible to microbiota-driven changes in a way that the CC is not, since myelination in the PFC has been previously speculated to be particularly plastic and susceptible to environmental factors (Tomlinson et al., 2016; Forbes and Gallo, 2017; Mount and Monje, 2017). This is further supported by observations of lack of differences in relative myelin-related protein expression in other brain regions, such as the frontal cortex (rather than the PFC), amygdala, cerebellum, hippocampus or striatum, as result of a GF environment, although ultrastructural myelin characteristics in these regions were notably not investigated (Hoban et al., 2016).

GF and SPF animals, regardless of genotype, showed comparable numbers of OPCs (PDGFRα positive cells) in both brain regions investigated. Interestingly, a reduction of mature oligodendrocytes (GST-pi positive cells) was observed in the PFC region alone in BACHD GF animals compared to BACHD SPF, whereas WT GF animals exhibited a reduction in the PFC and, to a lesser extent, in the CC, when compared to WT SPF control. Our results suggest that while OPC numbers may be unaffected, the proportion of OPCs that differentiate into mature oligodendrocytes may be lower in a GF environment. Alternatively, the decreased number of GST-pi positive cells in GF animals could suggest a potential increase in mature oligodendrocyte cell death driven by a GF status. Lower levels of myelin-related proteins in GF mice could be explained by a local decrease in mature oligodendrocytes. Notwithstanding, recent work has suggested that the extent of axonal myelination was independent of the number of newly formed and matured oligodendrocytes (Young et al., 2013). Thus, it is possible that the ability of mature oligodendrocytes to myelinate is compromised in a GF environment, leading to the observed decompaction of myelin lamellae, and alterations in the levels of myelin-related proteins.

Myelin breakdown disrupts white matter integrity in HD and has been identified as an early event in the pathology of the disease (Huang et al., 2015; Teo et al., 2016; Bourbon-Teles et al., 2017). Although mature oligodendrocytes were similarly affected by a GF environment in early manifest BACHD and WT animals, ultrastructural changes of myelinated axons were differentially expressed. These results suggest that the intrinsic pathological characteristics of BACHD mice may predispose these to myelination-related alterations due to environmental manipulations, such as gut microbiota depletion through GF housing.

A potential role for gut microbiota in HD pathology was inferred from differences identified in plasma metabolomic profiles of premanifest and early symptomatic HD subjects compared to healthy control (Rosas et al., 2015). Distinct gut microbiome-derived metabolites in the plasma of HD patients, therefore, suggested the mutant *HTT* may promote modifications in commensal gut bacteria. We did not observe any major global shifts in bacterial community, although there was a trend towards decreased richness and diversity at 6 months of age in BACHD mice (data not shown). Diversity at the microbial level can be challenging to interpret, with conflicting findings across studies and neurodegenerative disorders. For example, studies using Alzheimer disease (AD) mouse models have shown decreased (Minter et al., 2016) and increased (Harach et al., 2017) levels of microbiota diversity. In multiple sclerosis (MS), associated with alterations and disturbances of microbiota communities (Francis and Constantinescu, 2018), differences at the level of individual major taxa rather than shifts in diversity have been reported (Jangi et al., 2016; Berer et al., 2017; Cekanaviciute et al., 2017). Similar findings have been revealed in Parkinson disease (PD) patients (Scheperjans et al., 2015; Petrov et al., 2017). Therefore, the possibility arises that rather than lacking in bacterial diversity, affected individuals may harbour increased proinflammatory gut bacteria.

Further analysis of commensal bacteria at the family and genus level revealed only minor differences between BACHD and WT mice, where both 3 and 6 month old BACHD mice showed decreased abundance of *Prevotella* and *Bacteroides* at the genera level, part of the phylum Bacteroidetes. Consistent with our findings, *Prevotella* was previously reported to be decreased in the APP/PS1 transgenic mouse model of AD (Shen et al., 2017) and in the experimental autoimmune encephalomyelitis mouse, which recapitulates the main characteristic of MS (Mangalam et al., 2017). Moreover, microbiota profiling in humans also showed decreased *Prevotella* abundance in MS patients (Chen et al., 2016; Jangi et al., 2016), autistic children (Kang et al., 2013), and PD patients (Keshavarzian et al., 2015; Scheperjans et al., 2015), and was associated with increased gut permeability (Hasegawa et al., 2015). However, *Prevotella*, at high-levels, has also been linked to inflammation (Scher et al., 2013; Dillon et al., 2016) and, being a large genus, the role of its constituent members remains to be elucidated.

BACHD and WT animals in our facility were littermates and housed in the same cages in order to minimise confounding factors such as increased fighting or occurrences of cage-related stereotypies (Bailey et al., 2006). Thus, subtle effects may have been concealed by potential synchronisation of microbiota populations when cohousing wildtype and mutant littermates (Laukens et al., 2016). As a result, the functional significance of the modest differences in microbiota between BACHD and WT animals is difficult to interpret.

Whether manipulation of microbiota populations or microbiota derived metabolites in HD may dampen symptoms or ameliorate disease progression remains unclear. Interestingly, butyrate, one of the main short-chain fatty acids and a microbiota metabolite produced by *Prevotella* and *Bacteroides*, has been proposed to restore unbalanced gut microbiota, alter gene expression, and prevent neurodegeneration (Bourassa et al., 2016). Indeed, sodium butyrate and phenylbutyrate intake rescued histone acetylation, prevented neuronal death, and extended the lifespan of several HD mouse models (Ferrante et al., 2003; Gardian et al., 2005). Furthermore, thiamine, a by-product of *Bacteroidetes*, may also play a key role in the HD pathogenesis, where associated vitamin supplements benefit impaired energy metabolism in HD cells (Gruber-Bzura et al., 2017). Thus, further exploration of the therapeutic potential of microbiota modulation is warranted.

In summary, our findings indicate that myelination is differentially affected in BACHD and WT mice in the absence of microbiota. However, the lack of profound differences in bacterial populations between the genotypes may suggest that defects in myelin plasticity observed in the BACHD mouse model are intrinsic and not related to pre-existing differences in gut microbiota. Future studies are needed to examine the effects of microbiota manipulation and microbiota derived metabolites on HD-related neuropathological and behavioural manifestations, and to elucidate their utility as biomarkers or therapeutics in HD.

## ACKNOWLEDGMENTS

We thank members of the Pouladi lab for thoughtful discussions and comments. We thank Jing Ying Tan and Llanto Elma Faylon for mouse colony breeding, maintenance and genotyping; and Eveline Shevin and Amanda Chern for assistance with immunohistochemistry and immunoblotting procedures. M.A.P was supported by funds from the Agency for Science Technology and Research and the National University of Singapore. S.P. was supported by funds from LKC School of Medicine and SCELSE.

## ABBREVIATIONS

BACHD: bacterial artificial chromosome model of HD
CC: corpus callosum
GST-pi: Glutathione S-transferase
GF: germ-free
HD: Huntington disease
SPF: specific-pathogen free
*HTT*: huntingtin gene
MBP: myelin basic protein
OPC: oligodendrocyte precursor cells
PDGFRα: platelet derived growth factor receptor alpha
PFC: prefrontal cortex
PLP: proteolipid protein
WM: white matter
WT: wild-type.

